# Extending conventional TIRF microscopy to image single molecules in micromolar analyte backgrounds

**DOI:** 10.64898/2026.08.24.746893

**Authors:** R.C. Gentry, K.M. Leon Hernandez, R.L. Gonzalez, C.D. Kinz-Thompson

## Abstract

Weak, reversible interactions underpin biomolecular recognition, and single-molecule fluorescence (smF) imaging techniques can provide unprecedented insight into those biological processes. Unfortunately, such studies often require micromolar concentrations of fluorophore-labeled biomolecules, which is beyond the accessible range of conventional smF microscopies. Here, we describe a surface-functionalization method based on cloud-point polyethylene glycol (PEG) grafting that enables widefield smF microscopy measurements at micromolar concentrations without the use of nanophotonic devices. Using conventional total internal reflection fluorescence (TIRF) microscopy, we detected single-molecule fluorescence resonance energy transfer (smFRET) from surface-tethered, donor-labeled target molecules with up to 8 μM concentrations of freely diffusing, acceptor-labeled analyte molecules in the background—two orders of magnitude higher than typical studies in the literature. Weak, DNA-hybridization and protein-RNA binding equilibria were measured across micromolar range titrations. Together with advances in high-background data analysis, the robust method presented here enables kinetic and thermodynamic analyses of weak biomolecular interactions, especially those limited by nonspecific adsorption and high fluorescence backgrounds, using only standard smF instrumentation.

---

Single-molecule techniques are powerful experimental approaches, because they can resolve molecular heterogeneity and reveal mechanistic pathways obscured by ensemble averaging^1–3^. Of these techniques, widefield smF imaging is particularly effective, because it enables highly multiplexed data collection from thousands of fluorophore-labeled molecules^1^. One common modality investigates biomolecular recognition and binding between a surface-tethered, target molecule and a freely diffusing, fluorophore-labeled analyte. In this approach, the target-associated fluorescence signal must be distinguished from background fluorescence generated by unbound analytes. As the analyte concentration increases, however, this background fluorescence obscures the target-associated fluorescence signal, and ultimately compromises single-molecule detection—an effect known as the concentration barrier^4–7^.

Optical strategies that reduce the excitation volume can overcome the concentration barrier by extending the working concentration range of smF imaging. For example, TIRF microscopy confines the excitation to a shallow evanescent field at the imaging interface, reducing background fluorescence^1^. Alternatively, the excitation can be confined using nanophotonic devices, such as zero-mode waveguides (ZMWs), which are subwavelength apertures in an opaque metal film^8–10^. This confinement theoretically extends the working concentration range of ZMW-based smF imaging into the micromolar range^8–10^. Both conventional TIRF-based and ZMW-based imaging experiments can be performed using smFRET, a type of smF technique, as the fluorescence readout to further extend the working concentration range. In smFRET experiments, distance-dependent energy transfer from a donor fluorophore to an acceptor fluorophore provides additional selectivity, because acceptor emission predominantly arises only from analytes that are specifically associated with the target biomolecule^2,10^.

Despite the benefits of excitation-volume reduction strategies and FRET-based excitation selectivity, reported analyte concentrations in the literature for both TIRF-based and ZMW-based smF imaging studies remain well below the micromolar regime. Notably, ZMW-based smF studies of biomolecular mechanisms are typically performed with analyte concentrations in the tens-of-nanomolar range^11–15^, and only a handful of TIRF-based smFRET experiments have been reported that use concentrations above 100 nM^6,16^. Our groups, for example, have not previously published a TIRF-based smFRET binding experiment using more than 50 nM acceptor-labeled analyte^17,18^.

These reports reveal a gap between the accessible concentration range expected from excitation-volume reduction strategies and FRET-based excitation selectivity, and the range actually achieved in practice. For measurements of surface-tethered target molecules in a background of freely diffusing analytes, an important practical constraint is the nonspecific adsorption of analytes to the imaged surface^19–23^. In smF colocalization experiments, nonspecifically adsorbed analytes can artifactually appear to be bound to a target molecule or be mistaken for a surface-tethered target molecule^23–25^. More generally, excitation of even temporarily adsorbed analytes can generate a significant background signal that obscures the target-associated signal. Consequently, nonspecific adsorption can prevent surface-tethered smF experiments from reaching analyte concentrations high enough to efficiently measure weak, biomolecular interactions.

We recently reported two TIRF-based smFRET studies of eukaryotic protein synthesis in which fluorophore-labeled eukaryotic initiation factors (eIFs) exhibited essentially no noticeable nonspecific adsorption. In those studies, we only attempted to reach solution concentrations of 50 nM^18^ and 130 nM^26^, however, that notable level of performance was the result of a significant redevelopment and optimization of the surface chemistry used to functionalize the quartz microscopy flow cells. Here, we describe and validate that surface chemistry: an optimized, amine-free, direct PEG-silane grafting strategy under cloud-point conditions that efficiently suppresses nonspecific adsorption at micromolar analyte concentrations. Such concentrations still result in detrimental levels of background fluorescence signal, however, which challenge traditional smF data analysis approaches. By using autocorrelation function-based localization and frame-wise background estimation during smF data analysis to address the high background signal encountered at high analyte concentrations, we demonstrate that our approach enables conventional TIRF-based smFRET measurements in the micromolar regime. To demonstrate the effectiveness of this method, we readily quantify the kinetics and thermodynamics of weak, DNA-hybridization and protein-RNA-binding equilibria across micromolar analyte titrations using conventional TIRF microscopy instrumentation—without the need to employ nanophotonic devices.

## RESULTS

### The concentration barrier is higher than commonly assumed

For surface-tethered smF binding measurements, the concentration barrier is often heuristically estimated as the analyte concentration at which, on average, one freely diffusing analyte molecule is present within the observation region of a target molecule. For an effective TIRF observation volume of ∼5 attoliters, estimated using a diffraction-limited lateral observation area and an evanescent-field depth of ∼100 nm, this occupancy corresponds to an analyte concentration of ∼340 nM (see *Supporting Information)*. The fluorescence intensity generated by the freely diffusing analytes in this volume is the source of background fluorescence that ‘contaminates’ the target-associated fluorescence signal. However, the mean background fluorescence intensity value can be easily estimated and subtracted to yield the desired target-associated signal alone. This occupancy-based heuristic, therefore, does not capture the principal limitation at high analyte concentrations: fluctuations in the amount of background fluorescence, called shot noise, that prevent reliable discrimination between the presence and absence of the target-associated signal.

The concentration barrier is better understood by considering the limit imposed by photon shot noise on detecting a target-associated signal. In widefield, TIRF-based smF imaging systems, a camera records a time series of fluorescence images, which we refer to as a TIRF movie. In each frame of a TIRF movie, the camera-recorded pixel values can be used to estimate the number of photons, *z*, originating from the observation region of a target molecule. There are two cases that would give rise to a particular value of *z*: *N* number of freely diffusing analyte molecules contributing background fluorescence without a target-associated signal (Case A), or the same *N* analyte molecules together with a target-associated signal (Case B; Fig. 1a). The presence of a target-associated signal can be identified if the probability that *z* arose from the photon-count distribution for Case B is greater than for Case A. However, as *N* increases, the photon-count distributions increasingly overlap, making it more and more difficult to determine whether the target-associated signal is present. If the target-associated signal contributes an expected *ε* photons per frame, and each freely diffusing analyte contributes *ηε* photons per frame (where *η* is a fraction), then the expected photon counts for each case are *k_A_* = *Nηε* and *k_B_* = (*Nη* + 1)*ε*, respectively. Assuming enough photons can be detected to approximate Poisson distributed photon-counts as normal distributions with equal mean and variance, then the photon-count distributions are N(*z* ∣ *k_A_*, *k_A_*) and N(*z* ∣ *k_B_*, *k_B_*), respectively. Statistically, the target-associated signal then has the highest likelihood of being present if *z* is greater than the threshold value, *z_BA_*, at which point the two photon-count distributions have an equal probability (Fig. 1b). Thus, errors in identifying the presence of the target are determined by the false-negative identification probability that the *z* falls below *z_BA_*, which is 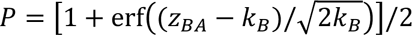 (Fig. 1b; see *Supporting Information* for a full derivation).

**Figure 1.**
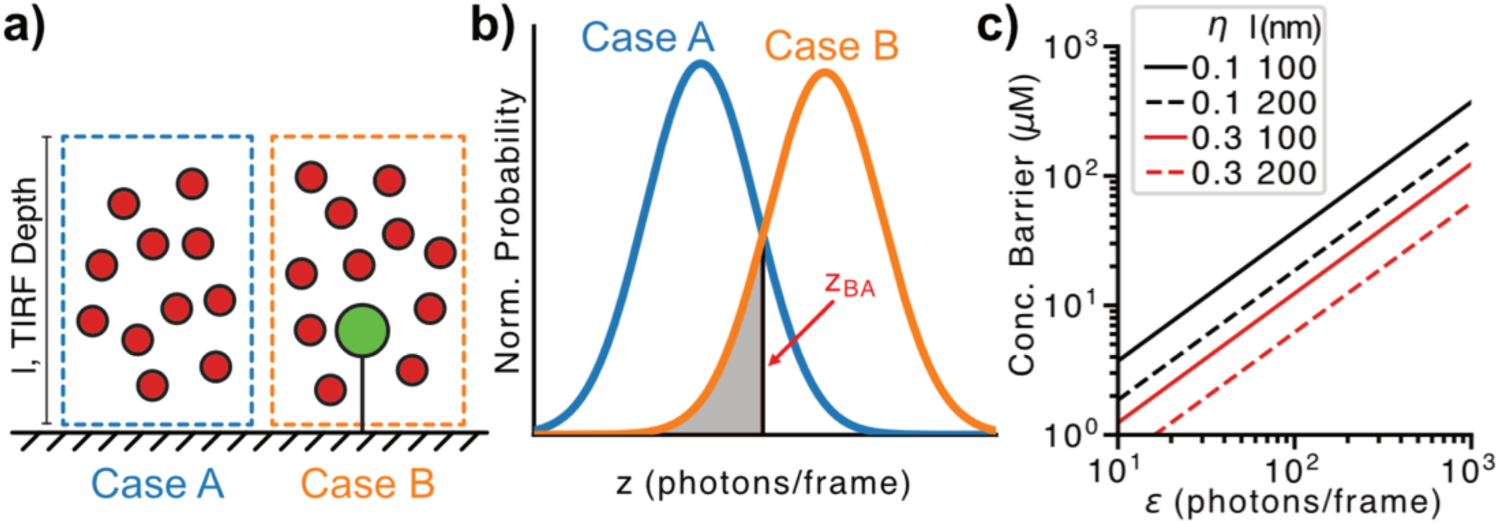
Shot-noise model of the concentration barrier. a) Schematic of two photon-detection cases within the observation region of a surface-tethered target molecule: *N* freely diffusing analyte molecules without a target-associated signal (Case A), or the same *N* analyte molecules together with the target-associated signal (Case B). Analyte molecules are shown in red, and the source of the target-associated signal is shown in green. b) Photon-count distributions for Case A (blue) and Case B (orange). The gray-shaded region below the threshold, *z_BA_*, represents the false-negative probability, *P*, that a target-associated signal is present but not detected. c) Estimated concentration barrier as a function of target-associated signal brightness, *ε*, for the indicated values of relative analyte brightness, *η*, and TIRF evanescent-field depth, *l*. The barrier occurs at a ∼10% false-negative probability.

The false-negative identification probability *P* increases from effectively zero to a maximum of 0.5 as the concentration of analyte molecules increases. The concentration barrier arises when *P* becomes significantly nonzero (*e.g.*, ∼0.1), which occurs when more than *N* ≥ *ε*/(2*πη*) analytes are within the observation region of a target molecule (Fig. 1c; see *Supporting Information)*. For a typical smFRET experiment, a bound, donor-excited acceptor molecule might contribute a conservative *ε* = 100 photons per frame, and *ηε* accounts for the direct excitation of each unbound acceptor-labeled analyte by the donor-excitation laser. With *η* = 0.1 estimated as the relative extinction coefficients of the acceptor and donor at the donor-excitation wavelength, the concentration barrier criterion is reached when *N* = 159, which corresponds to an analyte concentration of ∼54 μM at the assumed observation volume—two orders of magnitude above the conventional one-molecule estimate. At this concentration limit, the target-associated signal divided by the photon shot noise of the combined target-associated signal and analyte-generated background yields a shot-noise-limited signal-to-noise ratio (SNR) of ∼2.5.

The exact concentration limit depends on experimental factors such as the observation volume, fluorophore brightness, camera integration time, and relative brightness of the target-associated signal and freely diffusing analytes. Nonetheless, this model conservatively places the shot-noise-limited concentration barrier in the tens-of-micromolar range for TIRF-based smFRET experiments, and in the low-micromolar range for TIRF-based smF experiments without FRET-based selectivity (*η* = 1). Because only a few TIRF-based smFRET studies have even reached analyte concentrations in the hundreds-of-nanomolar range,^6,16^ this result suggests that factors other than shot noise from freely diffusing background analytes must commonly be determining the practical concentration limit.

### Amine-free PEG grafting under cloud-point conditions mitigates nonspecific adsorption

We hypothesized that incomplete functionalization of the microscope slide surface leads to significant nonspecific adsorption, and that these adsorbed molecules cause the gap between the theoretical and practical concentration limits in smF imaging. The conventional method for surface functionalization in smF imaging is a two-step process. First, the surface is aminosilanized to generate a layer of primary amines; second, those amines are coupled to *N*-hydroxysuccinimide ester-functionalized PEG to create a hydrophilic PEG layer that resists nonspecific biomolecular adsorption^19,20^. Incomplete coupling, however, can leave gaps in the PEG layer that promote nonspecific adsorption (Fig. 2a). Especially since positively charged amines induce a significant amount of nonspecific adsorption, secondary reactions to cap unreacted amines^23^ and/or blocking steps to coat the gaps^20^ are typically used to improve the surface quality. Alternative strategies avoid amine coupling altogether, such as direct silanization with PEG-silanes^22^, or by coating a hydrophobic dichlorodimethylsilane-functionalized surface with biotinylated bovine serum albumin and a surfactant^21^. To our knowledge, these alternative surface-functionalization strategies have not been shown to support conventional TIRF-based smF measurements with micromolar analyte concentrations.

**Figure 2.**
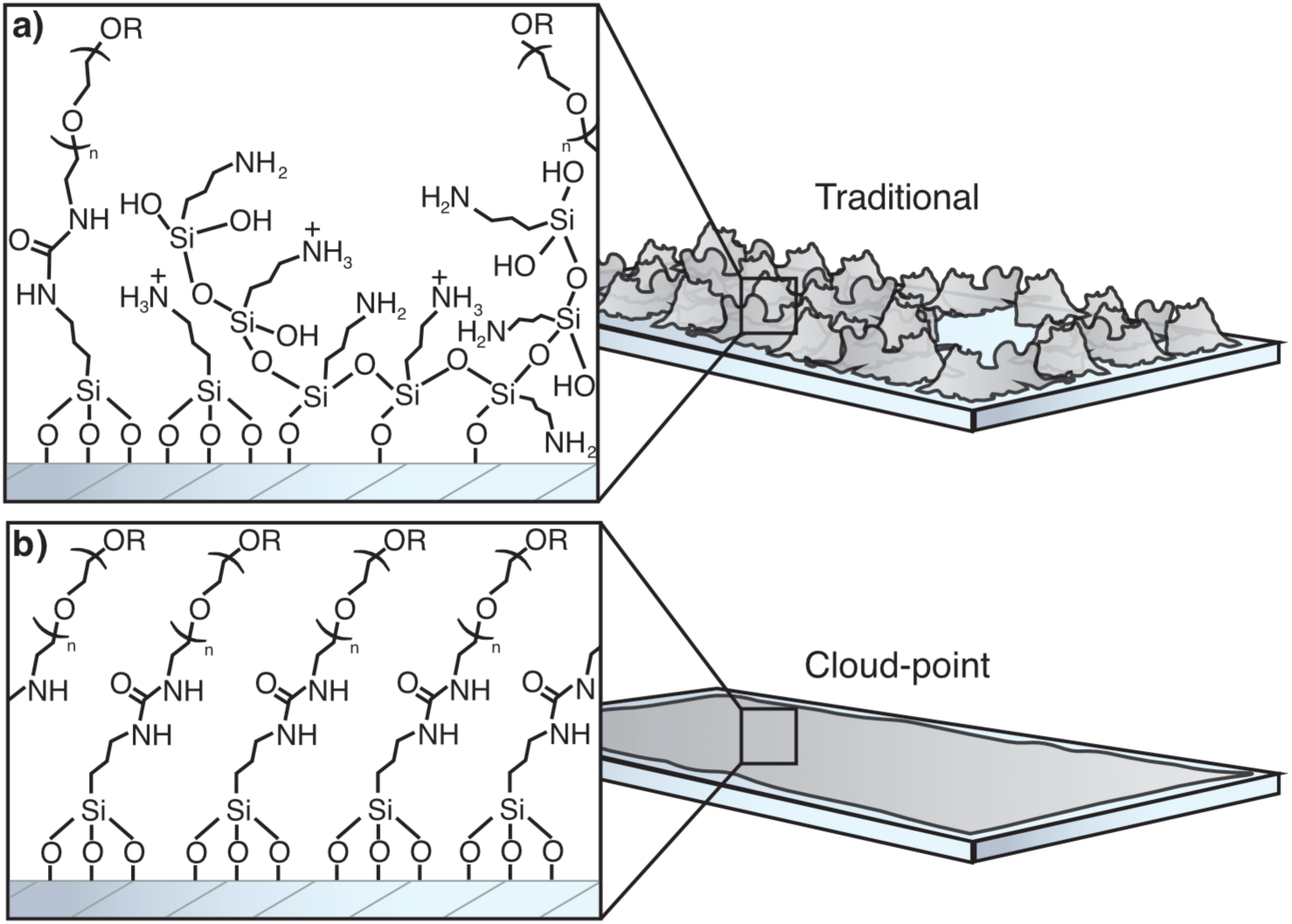
Comparison of conventional and cloud-point PEG surface functionalization. Molecular-scale (left) and microscale (right) schematics of quartz flow cell surfaces functionalized using the a) conventional aminosilane-PEG coupling method, or b) optimized, amine-free, direct grafting of PEG-silanes under cloud-point conditions. The conventional method couples N-hydroxysuccinimide ester-functionalized PEG to an aminosilanized surface, whereas the cloud-point method directly grafts methoxy-terminated PEG-silane and biotin-terminated PEG-silane onto the quartz surface.

To address these shortcomings, we have developed an amine-free strategy to improve surface functionalization by directly grafting PEG-silanes under cloud-point conditions, which promote the formation of a dense, homogeneous PEG layer (Fig. 2b). Direct silanization avoids the negative effects of amine-induced adsorption^22^, while still relying on the anti-fouling properties of PEG^27^. PEGylation was performed with a mixture of dilute amounts of biotin-terminated PEG-silane in methoxy-terminated PEG-silane to permit a controlled number of biotinylated target molecules to be tethered through a biotin-streptavidin-biotin bridge^20,22^. Finally, the functionalization efficacy was significantly improved by performing the silanization reaction under cloud-point conditions, which is known to reduce repulsion between PEG chains and induce a PEG conformation that promotes the formation of a more densely grafted PEG layer^23,28^

The cloud point of a PEG-silane solution depends on factors such as salt and PEG concentrations, PEG chain length, solvent, and temperature; thus, it should be empirically determined since it exhibits batch-to-batch variability. For reference, the cloud point of 5% *w/v* PEG-silane (MW=5 kDa) in 1% *v/v* acetic acid at room temperature is reached at ∼750 mM magnesium sulfate. Centrifugation of a cloud-point solution produces an aqueous biphasic system comprising a thin, upper phase that is enriched in PEG, and a thicker, salt-rich, lower phase that is depleted in PEG (Extended Data, Fig. 1). The lower phase is used for surface functionalization, because the reduced PEG concentration and reduction in interchain repulsion are expected to permit denser grafting^23,28^. The reduced PEG concentration necessitates a longer reaction time; however, catalytic amounts of acid were added to help limit silane self-condensation during this extended reaction period.

Under these cloud-point conditions, direct silanization using the lower phase yields surfaces that are so refractory to nonspecific adsorption that they allow TIRF-based smFRET experiments at micromolar concentrations (see below)^18,26^. Qualitative ‘sticking tests’ yielded negligible nonspecific adsorption after a one-minute incubation with a fluorophore-labeled protein, eIF4A (Extended Data, Fig. 2a). However, nonspecific adsorption was detectable at surface defects (*e.g.*, scratches) that are common to all quartz material, indicating that the procedure does not effectively functionalize these regions (Extended Data, Fig. 2b).

### TIRF-based smFRET imaging and analysis at micromolar concentrations

We next sought to test whether conventional TIRF-based smFRET imaging could finally be used to measure binding equilibria at micromolar analyte concentrations without the need for nanophotonic devices. At micromolar concentrations, however, background fluorescence is generally so high that it poses two substantial computational challenges to any smF analysis: (1) the resulting photon shot noise impedes the localization of surface-tethered target molecules in TIRF movies, and (2) it complicates the separation of the target-associated signal from the analyte-generated background. We handle the first challenge by identifying puncta in a pixel-wise autocorrelation function (ACF) image derived from each TIRF movie. At nonzero time lags, the ACF removes contributions from temporally uncorrelated noise sources, such as photon shot noise. Therefore, the ACF image enables shot-noise-free localizations under conditions in which puncta are otherwise difficult to identify in a conventional time-averaged image, which only reduces such noise (see *Supporting Information*). We handle the second challenge by simultaneously estimating the fluorescence intensity signal from a stationary, diffraction-limited spot (*i.e.*, at the location of a surface-tethered target molecule) and the local fluorescence background level at that spot in each frame by using maximum-likelihood estimation (see *Supporting Information*). Together, these ACF-based localization and frame-wise background-estimation methods are particularly effective for analyzing TIRF movies, and both have previously been used in studies from our groups^29,18,30,31,26^; however, they are crucial for experiments with high analyte concentrations that generate significant amounts of background fluorescence. We have now implemented both methods in highFRET, a free, open-source Python package that enables processing of high-concentration TIRF-based smF data on a standard laptop through command-line or graphical interfaces. For TIRF-based smFRET imaging, highFRET yields fluorescence intensity *versus* time trajectories that are used to calculate FRET efficiency (E_FRET_) *versus* time trajectories (E_FRET_ trajectories) that can be analyzed using tMAVEN software^32^. To verify the ability of these two approaches to extract E_FRET_ trajectories across a broad E_FRET_ range, we used highFRET to analyze an approximately equimolar mixture of four DNA-duplex smFRET standards with reported E_FRET_ values of 0.13, 0.40, 0.69, and 0.91^33^. The standards were imaged on a laboratory-built prism-based TIRF microscope. From 20 steady-state TIRF movies, highFRET identified and extracted 47,123 E_FRET_ trajectories that resolved the four expected E_FRET_ populations (Fig. 3a–d).

**Figure 3.**
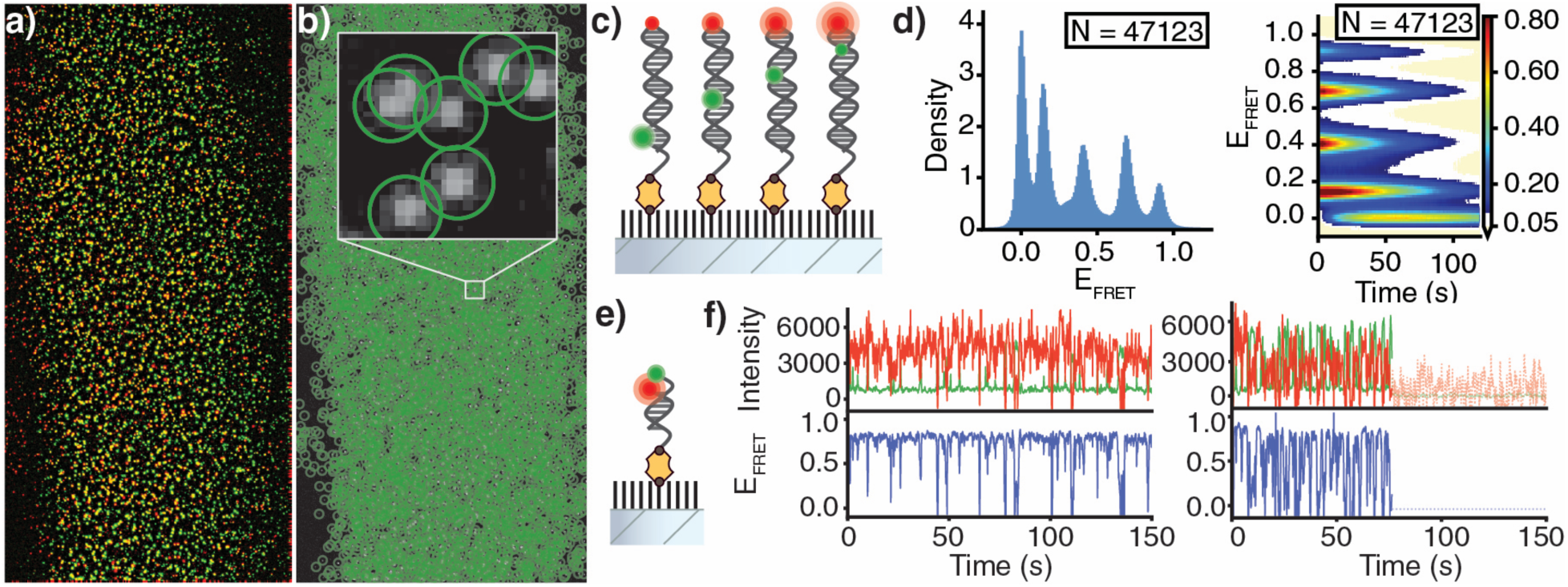
Analysis of smFRET data at micromolar analyte concentrations. a) Overlay of spatially aligned ACF images calculated from the donor (green) and acceptor (red) channels of a TIRF movie. b) Target-molecule locations identified as puncta in the donor ACF image. c) Schematic of the four double-stranded DNA (dsDNA) smFRET standards imaged simultaneously. d) Distribution of E_FRET_ (left) and E_FRET_ distribution as a function of time (right) for 47,123 E_FRET_ trajectories extracted from 20 TIRF movies of the dsDNA-standard mixture. The standards have reported E_FRET_ values of 0.13, 0.40, 0.69, and 0.91. (e) Schematic of the high-concentration DNA-hybridization smFRET experiment, in which Cy5-DNA analyte binds reversibly to a surface-tethered, Cy3-labeled DNA target molecule. (f) Representative E_FRET_ trajectories acquired at 4 μM Cy5-DNA using 25 mW excitation (left) and at 8 μM Cy5-DNA using 10 mW excitation (right). At 8 μM, the reduced excitation intensity decreased the target-associated signal, and the faster association kinetics increased temporal averaging, which together reduced the SNR.

We then tested the limits of TIRF-based smFRET imaging at high analyte concentrations by monitoring a weak, DNA-hybridization equilibrium (Fig. 3e). The surface-tethered target molecule was an 18-nucleotide single-stranded DNA bearing a 5′ biotin and a 3′ Cy3 fluorophore, which served as the donor. The analyte was a complementary 7-nucleotide DNA bearing a 3′ Cy5 fluorophore, which served as the acceptor (Cy5-DNA). The labeling sites were positioned 12 nucleotides apart in the duplex—a geometry predicted to produce a high E_FRET_ value (Online Methods). For this system, we recorded TIRF movies and extracted E_FRET_ trajectories that report on the binding equilibrium at Cy5-DNA concentrations up to 8 μM (Fig. 3f).

At 8 μM Cy5-DNA, however, the power of the excitation laser had to be reduced to avoid saturating the camera’s full-well capacity, thereby decreasing the SNR of the target-associated signal. Additionally, the binding dynamics were so rapid that they occurred within individual camera exposure periods, which yielded temporal averaging that also reduced the SNR^34^. Thus, on our conventional TIRF microscope, this new surface functionalization method turned the balance between detector dynamic range and sufficient excitation power into the limiting factor for TIRF-based smFRET imaging at analyte concentrations of approximately 5-10 μM.

### Measuring micromolar-affinity interactions at micromolar concentrations of analyte

With the concentration range accessible to TIRF-based smFRET imaging now expanded to micromolar concentrations, we next sought to quantify weak, biomolecular interactions that are difficult to measure without nanophotonic devices. Specifically, we examined: (1) a DNA-hybridization equilibrium with a room-temperature melting temperature, and (2) a protein-RNA binding equilibrium with a micromolar equilibrium dissociation constant (*K*_D_)^18^. Because the *K*_D_ for these systems is so large, accurate quantitation of the equilibria requires using analyte titrations that reach into the micromolar range.

Using the DNA-hybridization smFRET signal described above, we performed pre-steady-state binding experiments using solution exchange of Cy5-DNA (*i.e.*, similar to stopped-flow delivery; Fig. 4a). E_FRET_ trajectories were well resolved throughout a Cy5-DNA titration from 100 nM to 1 μM, and increasing the analyte concentration increased the fraction of target molecules occupying the hybridized state, which exhibits a high E_FRET_ value (Fig. 4b,c and Extended Data Fig. 3). Apparent association rates obtained from pre-steady-state and steady-state analyses increased linearly with analyte concentration, and dissociation rates were independent of analyte concentration (Extended Data Fig. 4). Some E_FRET_ trajectories transiently sampled a low E_FRET_ state, which might represent an encounter complex or another short-lived intermediate in the hybridization pathway^2^. Notably, the titration spanned the concentration at which half of the target molecules were hybridized. The *K*_D_ values obtained from the pre-steady-state and steady-state analysis agreed within uncertainty, demonstrating the ability to accurately extract both kinetic and thermodynamic parameters at analyte concentrations up to 1 μM (Extended Data Figs. 4,5).

**Figure 4.**
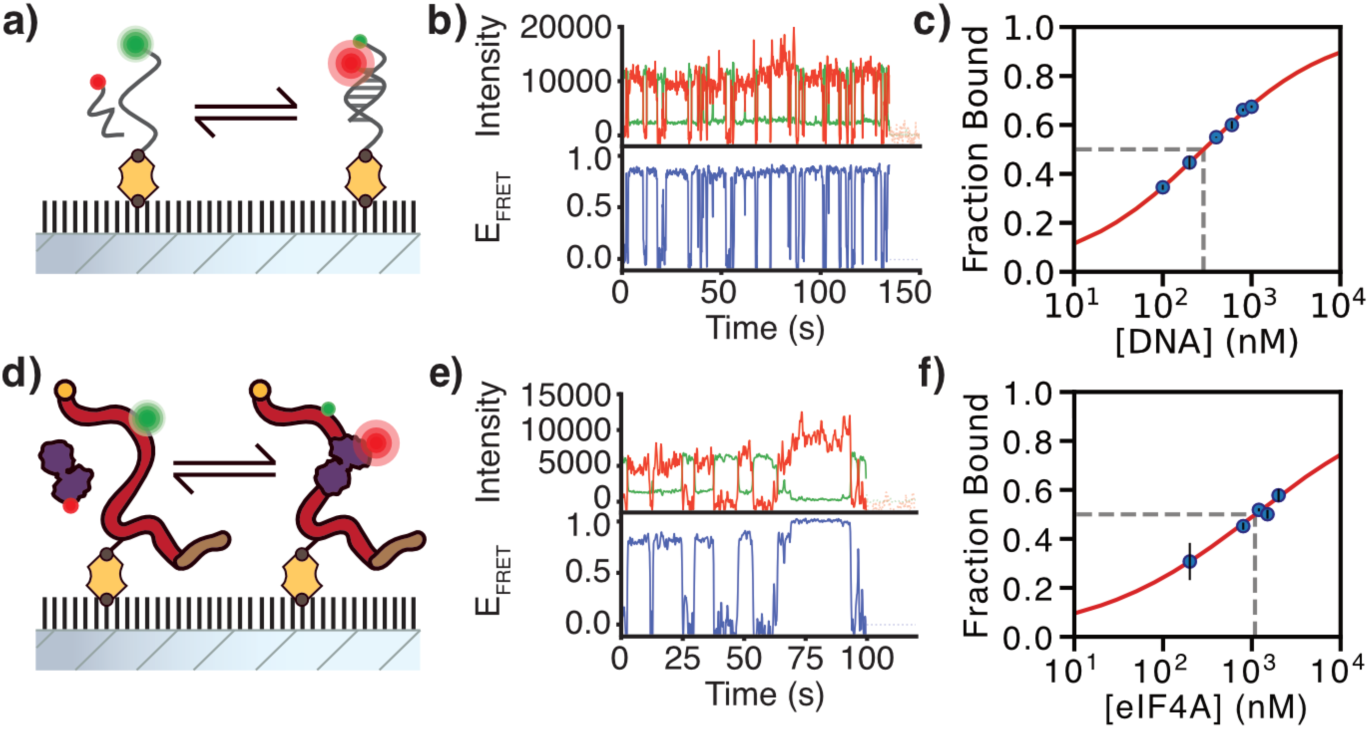
Micromolar smFRET titrations of DNA hybridization and eIF4A–RNA binding. a) Schematic of the DNA-hybridization experiment, in which a Cy5-DNA analyte binds reversibly to a surface-tethered, Cy3-labeled DNA target molecule. b) Representative E_FRET_ trajectory acquired at 1 μM Cy5-DNA. c) Cy5-DNA binding isotherm determined from smFRET measurements. Error bars are the standard deviation of two independent experiments. The solid line is fit to the Hill equation to guide the eye. d) Schematic of the protein-RNA binding experiment, in which a Cy5-eIF4A analyte binds reversibly to a surface-tethered, Cy3-labeled RNA target molecule. e) Representative E_FRET_ trajectory acquired at 2 μM total eIF4A, corresponding to 1.3 μM Cy5-labeled eIF4A based on the measured 65% labeling efficiency. f) eIF4A-binding isotherm determined from smFRET measurements. Error bars are the standard deviation of two independent experiments. The solid line is fit to the Hill equation to guide the eye.

We next tested a protein analyte, which we anecdotally note tend to be more adsorption-prone than nucleic acid analytes, by measuring the binding of Cy5-labeled eIF4A (Cy5-eIF4A) to a surface-tethered, Cy3-labeled RNA target (Fig. 4d). The target was a 24-nucleotide fragment from the 5′ untranslated region of the *Saccharomyces cerevisiae* rpl41a transcript^35^. This short fragment limits the number of RNA sites that eIF4A can occupy. Ensemble measurements place the *K*_D_ for eIF4A-RNA binding at approximately 4 μM^36–39^, a value that has been difficult to probe using conventional TIRF-based smFRET imaging.

In pre-steady-state experiments, Cy5-eIF4A was delivered by solution exchange at analyte concentrations ranging from 200 nM to 2 μM (Fig. 4e,f and Extended Data Fig. 6). Binding and dissociation events were resolved in individual E_FRET_ trajectories (Fig. 4e). Consistent with its weak affinity, eIF4A showed low RNA occupancy at 200 nM that became appreciable around 1 μM (Fig. 4f). Apparent association rates obtained from pre-steady-state and steady-state analyses also increased linearly with analyte concentration, although the association rate estimated at 200 nM was constrained by photobleaching of the Cy3-labeled RNA target; dissociation rates were independent of analyte concentration (Extended Data Fig. 4). The pre-steady-state and steady-state analyses yielded *K*_D_ values of 2.0 μM and 2.2 μM, respectively, close to the reported ensemble value 4 μM (Extended Data Fig. 5)^36–39^. These results demonstrate that surface functionalization *via* direct silanization under cloud-point conditions sufficiently mitigates nonspecific adsorption to enable the quantitative kinetic and thermodynamic analysis of weak biomolecular interactions using conventional TIRF-based smFRET imaging, with titrations that reach micromolar analyte concentrations.

## DISCUSSION

We developed an optimized, amine-free, cloud-point PEG-silane grafting strategy that produces microscopy flow-cell surfaces highly resistant to nonspecific adsorption of biomolecules at micromolar concentrations. The refractory nature of the treated surfaces, combined with ACF-based localization and frame-wise local background estimation, enabled conventional TIRF-based smFRET measurements that bridged the gap between the practical and theoretical concentration limits by allowing us to reach analyte concentrations up to 8 μM without using nanophotonic devices. However, this level of performance requires careful consideration when extracting target-associated signals from the high amount of analyte-generated fluorescence background.

Under conservative assumptions, our shot-noise model predicts that the theoretical TIRF-based smFRET concentration barrier is near 50 μM. This value is not universal; it depends on factors such as observation volume, fluorophore brightness, direct acceptor excitation, and microscope optics. Nonetheless, the gap between the predicted limit and the working concentrations reported in the literature indicates that surface adsorption and data processing, rather than photon shot noise alone, commonly determine the practical concentration range accessible in surface-tethered, TIRF-based smF experiments.

Although we focused on TIRF-based smFRET imaging, the surface chemistry presented here is independent of the fluorescence readout and can therefore benefit other modalities, including smF colocalization experiments^24,25^. However, because colocalization experiments lack the selectivity provided by FRET, their background-limited concentration range is lower. This surface chemistry approach can also be applied to borosilicate glass, as we have shown in interferometric scattering microscopy experiments^40^, and should also apply to silicon coated with an atomically thin SiO₂ layer for electrical sensing applications^41^. These applications suggest that this functionalization strategy may be useful whereever nonspecific adsorption limits measurements at solid-liquid interfaces.

Nonetheless, performing smF imaging at micromolar concentrations has important limitations. For example, success depends on the brightness and photophysics of the fluorophores, spectral crosstalk and spectral leakage, excitation depth, camera integration time, dynamic detector range, and the adsorption propensity of the analyte. Nonspecific adsorption propensity varies substantially among biomolecules, and surface functionalization will not be equally effective for all analytes. The procedure also does not passivate surface defects, such as scratches. Empirically, cloud-point conditions must be established for each PEG batch, and the prepared surfaces perform best when used within one to two days. Finally, no number of computational background-correction procedures can recover target-associated signals once photon shot noise makes the relevant photon-count distributions insufficiently distinguishable, or the detector becomes saturated.

Within these constraints, however, our groups now routinely use 200-300 nM Cy5-labeled analyte as the starting concentration for TIRF-based smFRET binding experiments performed on freshly prepared flow cells. Furthermore, this approach clearly extends conventional, surface-tethered, TIRF-based smFRET measurements into the micromolar regime. That capability enables kinetic and thermodynamic analysis of weak, biomolecular interactions using conventional TIRF instrumentation, and narrows the experimental gap between single-molecule and ensemble measurements.

## ONLINE METHODS

### Surface passivation

A detailed protocol is provided in the *Supporting Information*. Inlet holes were drilled into natural quartz microscope slides (1” × 3” × 1 mm; G. Finkenbeiner) using a 0.03-inch diamond burr drill bit (McMaster-Carr). The drilled slides were sonicated in ultrapure water for approximately 15 s to remove glass debris. Slides were immersed in 1% *w/v* Alconox for at least 2 hours, rinsed with ultrapure water and incubated in 1 M potassium hydroxide (KOH) for 2-3 hours. Slides and No. 1.5 borosilicate coverslips (24 mm × 30 mm; Fisher) were then sonicated sequentially in absolute ethanol for 20 minutes and 1 M KOH for 30 minutes, rinsed with ultrapure water, dried under 0.22-μm-filtered nitrogen or argon, and passed through a propane flame to remove organic impurities.

Methoxy-terminated PEG-silane (mPEG-silane; 5,000 Da; Laysan Bio) and biotin-terminated PEG-silane (biotin-PEG-silane; 5,000 Da; Laysan Bio) were combined at a 10,000:1 mass ratio. Specifically, 50 mg mPEG-silane was dissolved in 450 μL of 2% *v/v* aqueous acetic acid. Then, 20 mg of biotin-PEG-silane was dissolved in 1 mL of ethanol; 5 µL of that solution was then diluted into 1 mL of 2% *v/v* aqueous acetic acid; and finally 50 µL was added to the mPEG-silane solution. An equal volume of 1.5 M magnesium sulfate (MgSO_4_) was added to yield final concentrations of approximately 5% *w/v* total PEG-silane, 750 mM MgSO_4_, and 1% *v/v* acetic acid. The resulting cloudy mixture was vortexed, and centrifuged at >20,000 × g for 2.5 minutes at room temperature to form a thin, PEG-rich, upper phase and a thicker, PEG-depleted, lower phase. If two distinct phases were not formed, trials were conducted to identify a higher concentration of magnesium sulfate at which they formed.

For each slide-coverslip pair, 70 μL of the lower phase was applied between the cleaned surfaces. The assemblies were incubated for 20-22 hours at room temperature in a humid chamber. Slides and coverslips were then separated, rinsed thoroughly with ultrapure water, and dried under filtered nitrogen or argon. Flow cells were constructed using strips of double-sided tape and sealed with quick-drying epoxy. The assembled flow cells were used within one to two days.

### Flow-cell preparation and surface tethering of target molecules

Before imaging, each PEG-functionalized flow cell was flushed with 70 µL of 0.1% tween-20 resuspended in reaction buffer (see TIRF-Based smFRET microscopy). Next, the flow cell was washed twice with 70 µL of reaction buffer lacking tween-20. Then, 20 µL of 10 nM streptavidin was added to the flow cell, followed by two washes with 70 µL of reaction buffer. Finally, 70 µL of biotinylated substrate (50 pM – 500 pM) was introduced for 30 s before washing twice with 70 µL of reaction buffer. The concentration of biotinylated substrate concentration was adjusted until around 1200 spots were detectable in the donor channel. Target molecules were thereby tethered through a biotin-streptavidin-biotin bridge between biotin-PEG-silane on the surface layer and the biotinylated target molecule.

### DNA and RNA constructs

The dsDNA7, dsDNA10, dsDNA14 and dsDNA19 smFRET standards were prepared as described previously^33^ by hybridizing a biotin- and Cy5-labeled oligonucleotide (base) with the corresponding Cy3-labeled complementary oligonucleotide. High-performance liquid chromatography (HPLC)-purified oligonucleotides were purchased from Integrated DNA Technologies (IDT): 5′-Cy5-GGACTGCCGCCTGGGGAGCCGCACGACGACACGACAAAG-biotin-3′ (base); 5′-CGTGTCGTCGTGCGGCTCCCCAGGCG-Cy3-GCAGTCC-3′ (DNA7); 5′-CGTGTCGTCGTGCGGCTCCCCAG-Cy3-GCGGCAGTCC-3′ (DNA10); 5′-CGTGTCGTCGTGC GGCTCC-Cy3-CCAGGCGGCAGTCC-3′ (DNA14); and 5′-CGTGTCGTCGTGCG-Cy3-GCTCCC CAGGCGGCAGTCC-3′ (DNA19).

The DNA-hybridization equilibrium target oligonucleotide, 5′-biotin-TTTTTGGTGATGCGTGCT-Cy3-3′, and analyte oligonucleotide, 5′-GCATCAC-Cy5-3′, were purchased from IDT with HPLC purification. The target and analyte concentrations were determined using extinction coefficients at 280 nm of 166,000 M⁻¹cm⁻¹ and 75,900 M⁻¹cm⁻¹, respectively.

The Cy3-labeled, 24-nucleotide RNA target was prepared as described previously^18^. It was generated by hybridizing a biotinylated DNA-oligonucleotide (Biotin-CTTTCTCCACTTGGCTCTCAT) to the first 45 nucleotides of an rpl41a RNA (GGAGACCACATCGATTCAATCGAAATGAGAGCCAAGTGGAGAAAG). The RNA was capped using the vaccinia mRNA capping kit (NEB) according to the manufacturer’s instructions, and labeled with Cy3 at A14 using a terbium-assisted deoxyribozyme as previously described^18^. The RNA was purified using silica RNA clean up columns (Zymo), quantified using absorbance at 260 nm (474,300 M⁻¹cm⁻¹), and stored at −20 ℃. Before immobilization, the RNA was mixed with the bridging oligonucleotide at concentrations of 750 nM and 500 nM, respectively, in reaction buffer lacking magnesium, heated to 65 °C for 5 minutes, and then allowed to cool to room temperature. The annealed complex was then diluted 1:1,000 in reaction buffer before use.

### eIF4A expression, purification and labeling

A *Saccharomyces cerevisiae* eIF4A (TIF1) variant containing the C250A and T189C substitutions and an N-terminal hexahistidine tag followed by a tobacco etch virus (TEV) protease cleavage site was prepared as described previously^14,18,26^. Briefly, the protein was expressed in *Escherichia coli* BL21 RIPL (Agilent), purified by immobilized-metal-affinity chromatography, cleaved with TEV protease and further purified by size-exclusion chromatography into storage buffer containing 20 mM 4-(2-hydroxyethyl)-1-piperazineethanesulfonic acid (HEPES), pH 7.4, 100 mM potassium acetate (KOAc) and 10% *v/v* glycerol.

Purified protein was maintained with a four-fold molar excess of tris(2-carboxyethyl)phosphine (TCEP) as the sole reducing agent. eIF4A at 20 µM was reacted with a three-fold molar excess of sulfo-Cy5-maleimide (Lumiprobe) in 1% *v/v* dimethyl sulfoxide for 16 hours at 4 °C in storage buffer supplemented with TCEP. The reaction was terminated by desalting: unreacted maleimide-derivatized fluorophores were removed using a desalting column and a Superdex 75 10/300 GL column (Cytiva), both equilibrated in 20 mM HEPES, pH 7.4, 100 mM KOAc, and 10% *v/v* glycerol. Cy5-eIF4A was flash-frozen and stored at −80 °C in single-use aliquots.

The Cy5-eIF4A labeling efficiency and concentrations were determined by UV-visible absorption spectroscopy. The Cy5 concentration was determined using an extinction coefficient of 250,000 M⁻¹cm⁻¹ at 650 nm. The eIF4A concentration was determined using an extinction coefficient of 18,005 M⁻¹cm⁻¹ at 280 nm, and corrected for Cy5 background absorbance using a 4% correction factor from the absorbance peak at 650 nm to correct the absorbance peak at 280 nm. Using these concentrations, the Cy5-eIF4A labeling efficiency was 65%. Reported Cy5-eIF4A analyte concentrations refer to total eIF4A concentration, but association rates were corrected with a first-order correction by dividing by 0.65 to account for observing only labeled eIF4A.

### TIRF-based smFRET microscopy

DNA-hybridization and eIF4A-binding experiments were performed at a room temperature of 21 ± 1 ℃ in 20 mM HEPES, pH 7.4, 100 mM KOAc, and 3 mM magnesium acetate using a laboratory-built prism-based TIRF microscope described previously^18^. Samples were excited with a 532-nm diode-pumped solid-state laser (gem, Laser Quantum) and fluorescence was collected through a 60x, 1.2 NA water-immersion objective (Nikon). Donor and acceptor emission were separated using a Dual-View (Photometrics), and recorded using an iXon Ultra 888 electron-multiplying charge-coupled device camera (Andor) with 2x binning. The effective pixel size was 443 x 443 nm^2^, and the field of view was 113 x 227 μm^2^. TIRF movies were acquired with 300 EM gain, 1× pre-amp gain, 10.0 MHz readout mode, 0.60 μs vertical speed, and +1 clock voltage. DNA-hybridization experiments were performed using a 100 ms exposure time, and 25 mW laser power measured at the prism for Cy5-DNA concentrations less than 4 μM. Laser power was reduced to 18 mW for 4 μM Cy5-DNA experiments, and to 10 mW for both 6 μM and 8 μM Cy5-DNA experiments to avoid detector saturation. eIF4A-binding experiments were acquired using a 200 ms exposure time and 10 mW laser power. The microscope was controlled using Micro-Manager software^42^.

Samples were imaged in buffers that were supplemented with photostabilizing additives at 1% *w/v* glucose, 0.02 mg/mL glucose oxidase, 0.017 mg/mL catalase, 1 mM cyclooctatetraene, 1 mM nitrobenzoic acid, and 2 mM 6-hydroxy-2,5,7,8-tetramethylchroman-2-carboxylic acid (Trolox) aged to 2% oxidation by UV exposure. For pre-steady-state experiments, imaging was initiated approximately 1 s before an analyte-containing solution supplemented with the photostabilizing additives was delivered by solution exchange within the flow cell driven by a computer controlled syringe pump (J-Kem Scientific). Pre-steady-state experiments were performed in duplicate for Cy5-DNA concentrations of 100, 200, 400, 600, 800, and 1000 nM, and Cy5-eIF4A concentrations of 200, 800, 1200, 1500, and 2000 nM.

### DNA standards microscopy

dsDNA standards were oligomerized at a 10:1 molar ratio of the Cy3-labeled to biotinylated Cy5-labeled oligonucleotides in 10 mM Tris, pH 8.0, 50 mM NaCl, and 1 mM EDTA by thermal annealing from 100 to 4 ℃ over 6 hours^43^. This excess ensured that predominantly annealed dsDNA would only be tethered to the microscope slide surface. An approximately equimolar mixture of the four dsDNA standards was imaged after surface tethering 25 μL of 10 pM total dsDNA. Imaging was performed at a room temperature of 17 ℃ in 10 mM Tris, pH 7.4, and 50 mM KCl using a second, laboratory-built prism-based TIRF microscope. This microscope used an inverted RAMM body (Applied Scientific Instrumentation), a 514-nm Cobolt diode laser (Hübner), a 60x, 1.2 NA water-immersion objective (Olympus), an Optosplit II image splitter (Cairn Research) with Chroma filters ET575/50m-TRF, ZT561rdc-UF, and T635lpxr-UF2, and an ORCA-Fusion BT scientific complementary metal-oxide-semiconductor camera (Hamamatsu) with 1x binning. The effective pixel size was 108 x 108 nm^2^, and the field of view was 125 x 250 μm^2^. TIRF movies were acquired in fast-scan mode with a 10 ms exposure time and a 26 mW laser power measured at the prism. The microscope was controlled using Micro-Manager software^42^.

### TIRF movie and E_FRET_ trajectory analysis

TIRF movies were analyzed using vbscope. Target-molecule locations were identified from a pixel-wise ACF image calculated at a lag of τ = 2 frames. All local maxima pixels in the ACF image were considered as candidate puncta that might contain a molecule. Positive identification of a target molecule was made if the pixel value was greater than an acceptance threshold value of 0.15. Coordinates of target molecules were mapped between donor and acceptor fluorescence color channels using a 4^th^-order polynomial transform estimated from an image of a ZMW array localized in both color channels. Only molecules identified in both donor and acceptor channels were kept for analysis.

For each accepted target location and camera frame, maximum-likelihood estimation was used to separate target-associated donor and acceptor fluorescence intensities from the local fluorescence background. The point-spread-function model was taken as a two-dimensional Gaussian, integrated over the area of each pixel, using an 11×11 pixel region surrounding each local maximum spot. The background was modeled as a constant value for each spot region, with the value changing for each frame. Target-associated fluorescence intensity was estimated in both the donor and acceptor channel, I_D_ and I_A_, respectively. Intensity trajectories with an SNR less than 5 were removed before analysis. E_FRET_ was then calculated as I_A_/(I_D_+I_A_), and a Chung-Kennedy filter was applied with a sensitivity exponent of 2, the recommended filter windows lengths of 2, 4, 6, 8, and 10, and a perturbation window width of 10. E_FRET_ trajectories were processed and modeled in tMAVEN^32^, and retained if they exhibited single-step photobleaching or a single-step photoblinking event, or if they displayed similar photophysical and intensity profiles to trajectories that did photobleach.

After photobleaching points were manually set in tMAVEN, each E_FRET_ trajectory was independently modeled with a three-state hidden Markov model using the vbFRET algorithm, and emission states across all trajectories were clustered using K-means. The two states with the highest E_FRET_ values were assigned to the bound state, and the remaining state was assigned to the unbound state. Steady-state rate constants were calculated using a survival analysis of the dwell-time distributions for the bound and unbound states^18^. Reported rates were the population average of the rate constants from non-linear least squares fitting of the dwell-time distributions to biexponential decays. Pre-steady-state association rate constants were also calculated using survival analysis by mono-exponential fitting of the first-unbound lifetime distribution. For each analyte concentration, the bound fraction was calculated as *f* = (1 + *k_d_*/*k_a_*)^−1^, where *k_a_* is the apparent association rate constant and *k_d_* is the dissociation rate constant. The bimolecular association rate constant was calculated from linear regression of the apparent association rate constants, corrected for labeling efficiency, *versus* the analyte concentration. The apparent association rate constant at 200 nM eIF4A was limited by photobleaching, so it was excluded from the bimolecular association rate constant calculation. The *K*_D_ was obtained as the average dissociation rate constant divided by the bimolecular association rate constant.

### Nonspecific-adsorption measurements

Flow cells were prepared as described above except that streptavidin and biotinylated target molecules were omitted. Each flow cell was incubated with 1, 10, or 100 nM Cy3-eIF4A for 1 min at a room temperature of 21 ± 1 ℃. Before imaging, the unbound protein was washed out, and then TIRF movies of 200 frames were collected.

## DATA AVAILABILITY

Raw TIRF movies and E_FRET_ trajectories selected for further analyses are available in a Zenodo repository (DOI: 10.5281/zenodo.22075398; public upon publication).

## CODE AVAILABILITY

vbscope software is available at https://github.com/GonzalezBiophysicsLab/vbscope-paper^29^. highFRET software is available at https://github.com/ckinzthompson/highfret, and the archived version used in this study is available in a Zenodo repository (DOI: 10.5281/zenodo.22075398; public upon publication). tMAVEN software is available at https://github.com/GonzalezBiophysicsLab/tmaven^32^.

## ACKNOWLEDGEMENTS

R.C.G. was supported by the NSF Graduate research fellowship program (DGE1644869), and the Helen Hay Whitney Foundation Research Fellowship. K.M.L. was supported by the NSF Garden State Louis Stokes Alliance for Minority Participation (GS-LSAMP) Bridges to the Doctorate program (EES1905142), and the NIH Graduate Research Training Initiative for Student Enhancement program (5T32GM140951). R.L.G. acknowledges support from the NIH through grants R35-GM153724, R01-CA277727, and R01-DK135088. C.D.K. acknowledges support from the NSF through grants CHE2137630 and CHE2442804. We thank the staff at the Columbia Precision Biomolecular Characterization Facility (PBCF) for access to molecular biology and biophysical instrumentation.

## AUTHOR CONTRIBUTIONS

C.D.K. conceived the project, and C.D.K. and R.L.G. supervised the project together. All authors wrote and edited the manuscript. R.C.G. optimized surface functionalization chemistry. R.C.G. and K.M.L. prepared DNA samples; R.C.G. prepared eIF4A and RNA samples. R.C.G. and K.M.L. performed TIRF-based smFRET experiments, and analyzed smFRET data.

