## Extended Data for "Extending conventional TIRF microscopy to image single molecules in micromolar analyte backgrounds"

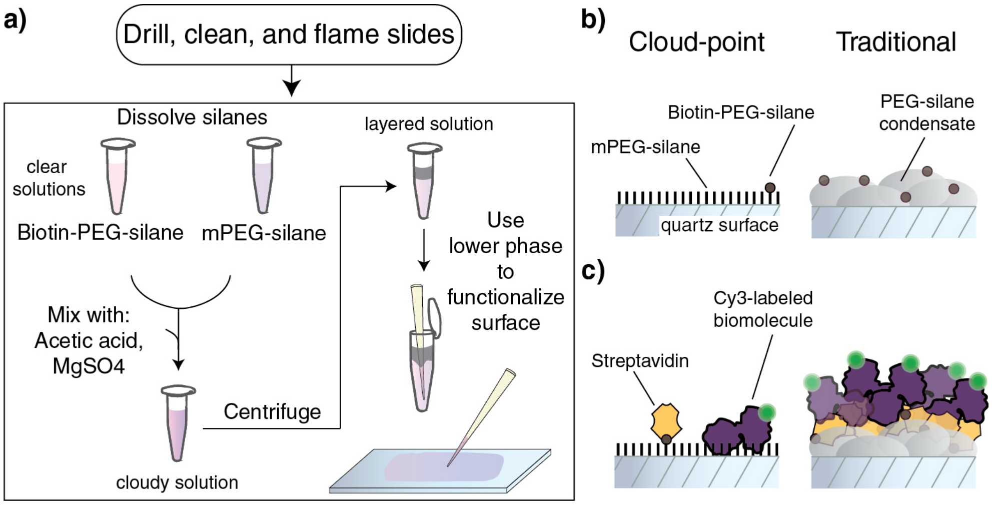


**Extended Data, Figure 1. Improved cloud-point functionalization procedure reduces nonspecific adsorption.** a) Schematic diagram of surface functionalization procedure. The cloud point depends on the PEG molecular weight, pH, temperature, solvent, and the identity and concentration of the salt. The final cloud-point solution should be made at the desired mPEG-silane:biotin-PEG-silane ratio. b) Cartoon of the distribution of PEG on a quartz surface functionalized under cloud-point (left) or traditional (right) functionalization approaches. c) Cartoon of the distribution of nonspecifically adsorbed fluorophore-labeled biomolecules bound to streptavidin-bound surfaces functionalized under cloud-point (left) or traditional (right) approaches.

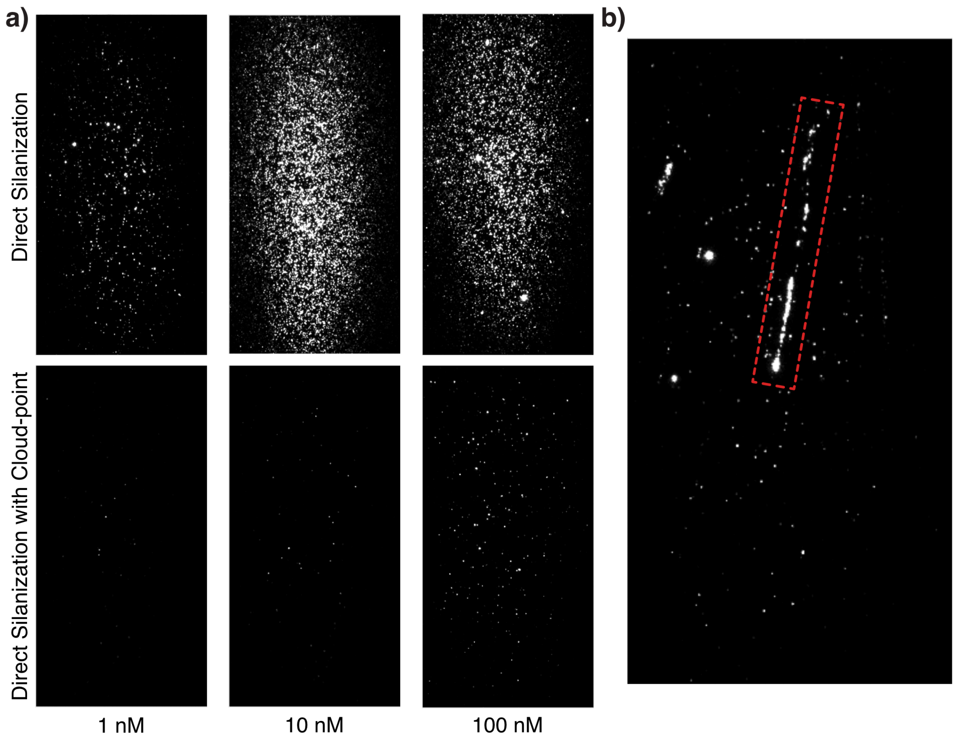


**Extended Data, Figure 2. Cloud-point functionalization protects microscope slide surfaces from nonspecific adsorption.** a) Representative images of a roughly 28,000 $\mu m^2$ field of view taken after incubating microscope slides functionalized using direct silane grafting either with or without the cloud-point approach as a function of increasing concentration of Cy3-labeled eIF4A. b) Representative image of nonspecific adsorption of Cy3-eIF4A to a surface defect caused by a scratch (red box) on a cloud-point functionalized surface without blocking protein.

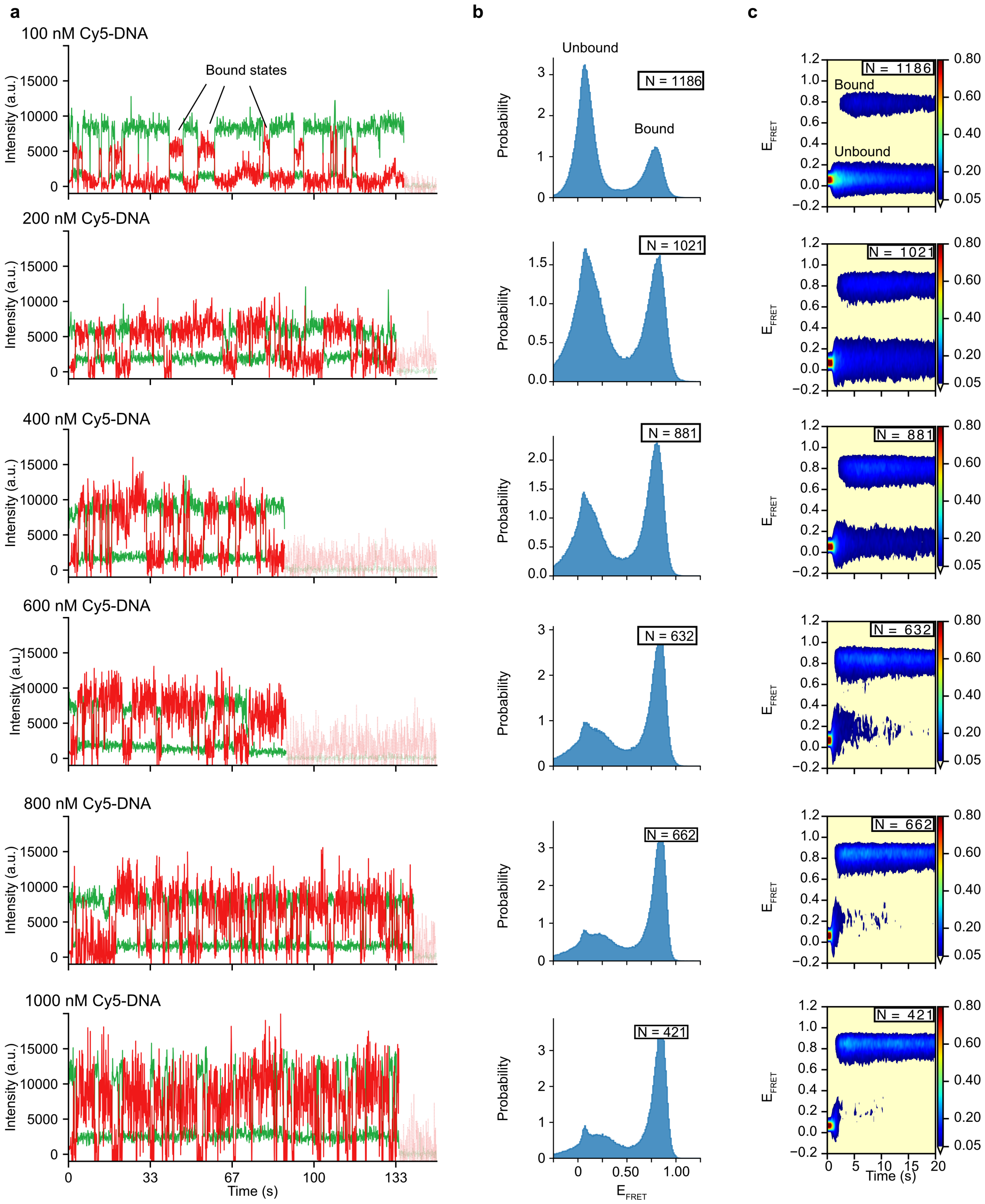


**Extended Data, Figure 3. Pre-steady state smFRET measurements of DNA hybridization at elevated concentrations for a titration series of 100 nM to 1 $\mu$M Cy5-DNA.** a) Representative example fluorescence intensity *versus* time trajectories of donor (green) and acceptor (red) fluorescence. Faded sections denote post-photobleaching measurements. b) Normalized histograms of $E_{FRET}$ distributions. c) Time-dependent histograms of $E_{FRET}$ distributions with colorbar mapping normalized to the maximum value in the histogram. *N* is the number of time trajectories in each histogram.

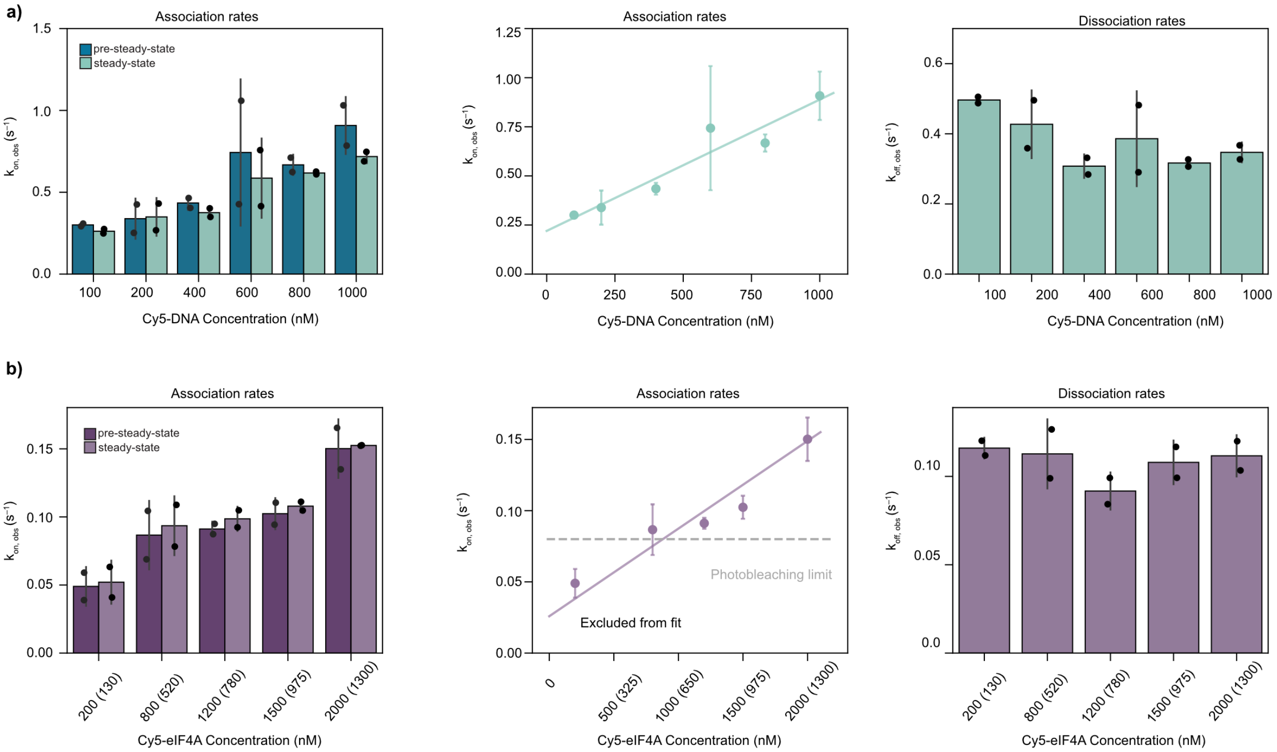


**Extended Data, Figure 4. Kinetic analysis of smFRET measurements.** a) Apparent association rate constants and dissociation rate constants for DNA hybridization as a function of Cy5-DNA concentration. b) Apparent association rate constants and dissociation rate constants for eIF4A-RNA binding as a function of Cy5-eIF4A. Concentrations are total eIF4A concentration, with Cy5-eIF4A concentration in parentheses. Linear regression (colored line) of the apparent association rates was used to obtain the bimolecular association rate constant. The 200 nM eIF4A concentration point was excluded from the linear regression due to limitations from photobleaching. Black points are results of individual experiments; bars and colored points are the average values; error bars are the standard deviation of two replicates.

| | $k_{on}$ ($M^{-1}$ $s^{-1}$) | | $k_{off}$ ($s^{-1}$) |
|---|---|---|---|
| Equilibrium | Pre-steady-state | Steady-state | Steady-state |
| DNA hybridization | 670000 ± 100000 | 500000 ± 50000 | 0.38 ± 0.09 |
| eIF4A-RNA binding | 53000 ± 16000 | 49000 ± 14000 | 0.11 ± 0.01 |

| | Affinity $K_D$ (μM) | |
|---|---|---|
| Equilibrium | Pre-steady-state | Steady-state |
| DNA hybridization | 0.57 ± 0.16 | 0.75 ± 0.20 |
| eIF4A-RNA binding | 2.0 ± 0.7 | 2.2 ± 0.7 |

**Extended Data, Figure 5. Summary of kinetic parameters determined by smFRET experiments for DNA hybridization and eIF4A-RNA binding.** Rate constants were determined as the slope from a logistic regression ($k_{on}$) or as the average ($k_{off}$) of data in Extended Data, Figure 4. Uncertainty is fitting error ($k_{on}$) or standard deviation ($k_{off}$). Equilibrium dissociation constants are calculated as the ratio of the off-rate constant to either the pre-steady-state or steady-state on-rate constant.

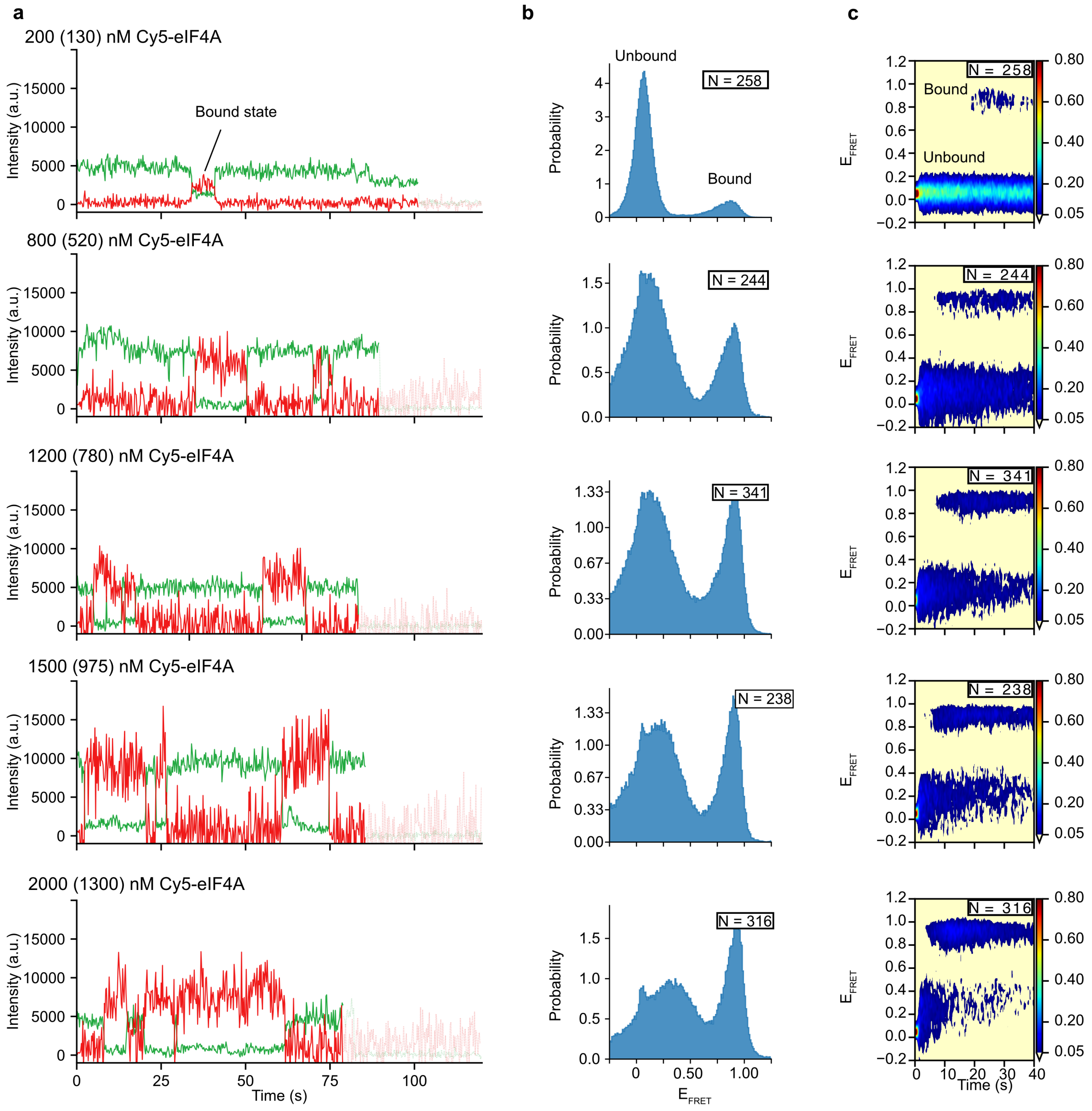


**Extended Data, Figure 6. Pre-steady state smFRET measurements of eIF4A-RNA binding at elevated concentrations for a titration series of 200 nM to 2 μM eIF4A.** Given concentrations are total eIF4A concentration with fluorophore-labeled Cy5-eIF4A concentration in parentheses. a) Representative example fluorescence intensity *versus* time trajectories of donor (green) and acceptor (red) fluorescence. Faded sections denote post-photobleaching measurements. b) Normalized histograms of $E_{FRET}$ distributions. c) Time-dependent histograms of $E_{FRET}$ distributions with colorbar mapping normalized to the maximum value in the histogram. *N* is the number of time trajectories in each histogram.
