## Supporting Information for "Extending conventional TIRF microscopy to image single molecules in micromolar analyte backgrounds"

### TABLE OF CONTENTS

1. Concentration of one molecule in a diffraction-limited volume
2. Shot-noise-limited concentration barrier derivation
3. Widefield single-molecule fluorescence image processing
4. Detailed cloud-point PEG-silane surface-functionalization protocol
5. References

### 1. Concentration of one molecule in a diffraction-limited volume

In the XY plane, the Abbe diffraction limit is $l_{xy} = \lambda/(2NA)$ for an imaging wavelength, $\lambda$, imaged through a lens with a numerical aperture, $NA$. For $\lambda$ = 532 nm light with a $NA$ = 1.2 objective, $l_{xy}$ = 221.7 nm. A square area, like that imaged onto a camera pixel, of this dimension has an area of $A_{sq}$ = 49,151 nm$^2$. Assuming that the evanescent field of a total internal reflection fluorescence (TIRF) microscope is just a hard cutoff at $l_z$ = 100 nm deep, then the approximate volume imaged onto this square area is $V \approx A_{sq} \cdot l_z = 4.9 \times 10^6\ nm^3$, which, when converted to liters is $V = 4.9 \times 10^{-18}\ L = 4.9\ aL$. Thus, the concentration in this region when it is occupied by one molecule is $C = 1/(V \cdot N_A) = 3.38 \times 10^{-7} M = 338\ \mu M$.

### 2. Shot-noise-limited concentration barrier derivation

To define a shot-noise-limited concentration barrier, consider two competing models for the number of observed photons, $z$, measured within the observation region of a target molecule during one camera frame: (A) the region only contains $N$ freely diffusing analyte molecules that generate background fluorescence, or (B) the region contains the same $N$ analyte molecules together with a target molecule that generates a target-associated fluorescence signal. The target-associated signal contributes an expected $\epsilon$ photons per frame, whereas each analyte contributes $\eta\epsilon$ photons per frame, where $\eta$ is a dimensionless multiplier describing the brightness of each analyte relative to the target-associated signal. In both cases, $z$ is distributed according to a Poisson distribution with rate parameter $k_A = N\eta\epsilon$, or $k_B = (N\eta + 1)\epsilon$, for cases A and B, respectively.

When $k_A$ is sufficiently large (*i.e.*, $k_A > 100$), both Poisson distributions can be approximated by heteroscedastic normal distributions,

$$\mathcal{N}(z|\mu = k_i, \sigma^2 = k_i u) = \frac{1}{\sqrt{2\pi k_i u}} e^{-\frac{(z-k_i)^2}{2k_i u}},$$

where $\mu$ and $\sigma^2$ are the mean and variance, respectively; $i \in \{A, B\}$; and $u$ is a constant of unit magnitude with the same units as $k_i$ to help to maintain dimensional consistency.

We can then define a threshold, $z_{BA}$, such that an observation of $z$ is more likely to correspond to case A when $z < z_{BA}$, or case B when $z > z_{BA}$. This threshold occurs at the crossover point where

$$\mathcal{N}(z_{BA}|k_A, k_A u) = \mathcal{N}(z_{BA}|k_B, k_B u).$$

This equality is satisfied at the roots of the quadratic equation

$$0 = (k_A u - k_B u) z_{BA}^2 + 0 \cdot z_{BA}^1 + \left(k_A k_B^2 u - k_A^2 k_B u + k_A k_B u^2 \ln \frac{k_B}{k_A}\right) z_{BA}^0,$$

which, under the conditions described above, where $k_A = N\eta\epsilon$ and $k_B = (N\eta + 1)\epsilon$, has only one meaningful solution of

$$z_{BA} = \sqrt{N\eta\epsilon(N\eta + 1)\left(\epsilon + u \ln \frac{N\eta + 1}{N\eta}\right)}.$$

When $\epsilon$ is sufficiently large that the logarithmic term is negligible, this expression simplifies to

$$z_{BA} \approx \sqrt{N\eta\epsilon(N\eta + 1)\epsilon}.$$

The concentration barrier emerges when $N$ becomes so large that it is not readily apparent whether a measurement of $z$ photons corresponds to case A or case B. This occurs because the photon-count distributions increasingly overlap as $N$ increases, which causes an observation generated under Case B to become marginally more and more attributable to Case A. The criterion described above where a measurement of $z$ photons is more likely to have originated from a region lacking a target molecule if $z < z_{BA}$ or more likely to have originated from a region containing a target molecule if $z > z_{BA}$ is equivalent to a maximum likelihood probabilistic inference calculation (*i.e.*, a comparison of whether $\mathcal{N}(z|k_A, k_A u) > \mathcal{N}(z|k_B, k_B u)$, or *vice versa*). Within this framework, the probability that we are unable to identify a region containing a target molecule is the false-negative identification probability

$$P = \int_{-\infty}^{z_{BA}} \mathcal{N}(z|k_B, k_B u)\, dz = \frac{1}{2}\left[1 + \operatorname{erf}\left(\frac{z_{BA} - k_B}{\sqrt{2k_B u}}\right)\right].$$

While this probability depends explicitly on $k_B$, it also depends on $k_A$ through the value of $z_{BA}$. Under nearly all experimental conditions, $\epsilon \gg 1$; so, in terms of molecular parameters, this probability is

$$P \approx \frac{1}{2}\left[1 + \operatorname{erf}\left(\frac{\sqrt{N\eta\epsilon} - \sqrt{(N\eta + 1)\epsilon}}{\sqrt{2u}}\right)\right].$$

This probability has a maximum of 0.5, which occurs when the photon-count distributions completely overlap (*i.e.*, when $\mathcal{N}(z|k_A, k_A u) \approx \mathcal{N}(z|k_B, k_B u)$), because under those conditions it is equally likely that any value of $z$ came from either case A or B. On the other side, this probability continuously approaches zero. However, like any probability, it is never exactly zero—just effectively zero. Since the concentration barrier emerges at the point where this probability becomes non-zero, but it is technically non-zero everywhere, we operationally define the concentration barrier as the point where the probability becomes *substantially* non-zero.

A reasonable choice is then the point where the first-order expansion of $P$ becomes zero, which occurs at about (Fig. S1). Truncated at the first-order term, the expansion of the error function is

$$\mathrm{erf}(x) = \frac{2}{\sqrt{\pi}}\left(x - \frac{1}{3}x^3 \dots\right) \approx \frac{2}{\sqrt{\pi}}x.$$

Using this in the expression for the false-negative identification probability, the resulting first-order approximation of $P$ is

$$P \approx \frac{1}{2} + \frac{1}{\sqrt{2\pi u}}\left(\sqrt{N\eta\epsilon} - \sqrt{(N\eta + 1)\epsilon}\right).$$

This expression for $P$ equals zero when

$$\sqrt{\frac{\pi u}{2\epsilon}} = \sqrt{N\eta + 1} - \sqrt{N\eta},$$

which can be solved by noting that

$$A = \sqrt{x + 1} - \sqrt{x} \quad \rightarrow \quad x = \frac{(A^2 - 1)^2}{4A^2}.$$

Solving for $N$, we obtain the number of background analyte molecules at which the false-negative identification becomes substantially non-zero. That value is where the value of $N$ at which concentration barrier emerges,

$$N_{CB} = \frac{(\pi u - 2\epsilon)^2}{8\pi\eta\epsilon u}.$$

This expression is dimensionless, as expected. Notably, for most experimental situations $\epsilon \gg \pi u/2$, which simplifies the expression for $N_{CB}$ to

$$N_{CB} \approx \frac{\epsilon}{2\pi\eta u}.$$

Thus, the shot-noise-limited concentration barrier occurs when

$$N \geq N_{CB} \approx \frac{\epsilon}{2\pi\eta u}.$$

Since $N_{CB}$ marks the point when the first-order approximation of the false-negative identification probability equals zero, using the full expression for $P$ and calculating $P(N_{CB})$ yields the probability that is considered to be "substantially" non-zero. Notably, this expression depends only on $\epsilon$, and as a function of $\epsilon$, it asymptotes to a value of 0.105 (Fig. S1). Thus, an operative definition of the concentration barrier is that it occurs when the false-negative identification probability becomes ~10%.

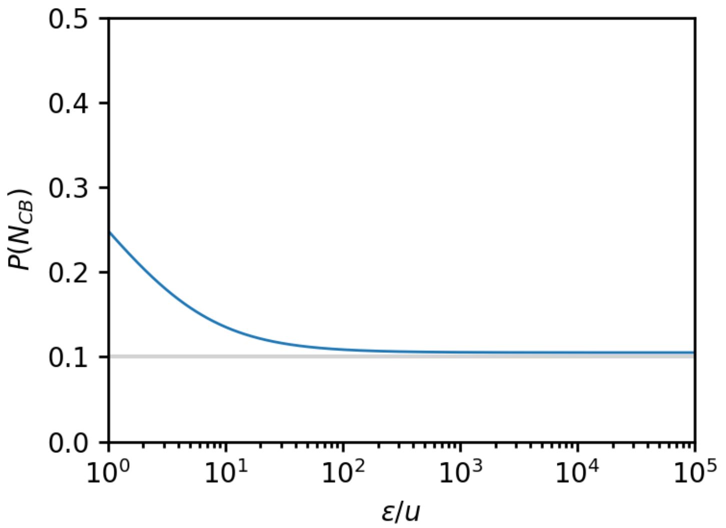


Figure S1. False-negative identification probability at the concentration barrier as a function of target-associated signal. The exact probability (blue) is evaluated at the point where the first-order approximation of the probability becomes zero. The grey line shows the asymptotic value of 0.105.

**3. Widefield single-molecule fluorescence image processing**

In the widefield, TIRF-based single-molecule fluorescence (smF) experiments considered here, a camera acquires fluorescence images at regular time intervals to generate a TIRF movie. Analysis of such surface-tethered TIRF-based smF measurements, including TIRF-based single-molecule fluorescence resonance energy transfer (smFRET) measurements, requires extracting fluorescence intensity *versus* time trajectories (intensity trajectories) for each target molecule. Intensity-trajectory extraction comprises two principal steps: (1) localizing each target molecule, and then (2) estimating the target-associated fluorescence intensity in each movie frame. We describe these steps below.

*Step 1: Localizing target molecules.* Surface-tethered target molecules can be localized by applying a detection criterion to each local maximum pixel in a fluorescence image to determine whether a diffraction-limited, target-associated fluorescence signal is present at that location. One conventional approach is to estimate the time-averaged image of the field of view using some portion of a TIRF movie:

$$\bar{I}_{xy} = \frac{1}{T}\sum_{t=t_1}^{t_T} I_{xy}(t)$$

where $I_{xy}(t)$ is the fluorescence intensity recorded at pixel $xy$ in time $t$, the $t_i$ are frames in the TIRF movie, and $T$ is the number of frames included in the average. The time-averaged image is often calculated from, for example, the first 20 frames of a movie or from control frames acquired while the field of view is illuminated with a different laser. Target molecules are then located by identifying local maxima pixels in the averaged image, and comparing the $\bar{I}_{xy}$ of each local maximum to an intensity threshold value, $I_0$. A target molecule is determined to be present at pixel $xy$ when $\bar{I}_{xy} \geq I_0$. Time averaging reduces, but does not eliminate, the effects on this estimate of noise such as photon shot noise or camera noise. Those noise sources impede localization, especially when the background fluorescence is high.

An alternative approach to target molecule localization is to identify them in a normalized, two-point autocorrelation function (ACF) image. For each pixel in a TIRF movie, the normalized ACF is calculated as

$$G_{xy}(\tau) \equiv \frac{\mathrm{ACF}_{xy}(\tau)}{\mathrm{ACF}_{xy}(\tau = 0)}$$

where the raw ACF is estimated as

$$\mathrm{ACF}_{xy}(\tau) \approx \frac{1}{T - \tau} \cdot \sum_{t=1}^{T-\tau} \left[ I_{xy}(t) \cdot I_{xy}(t + \tau) \right]$$

and $\tau$ is a time lag (*e.g.*, $\tau = 2$ camera exposures). The centered ACF can also be used by subtracting an estimate of the mean from the signal intensity, $\delta I_{xy}(t) \equiv I_{xy}(t) - \bar{I}_{xy}(t)$, before calculating the ACF. More details on using ACFs for molecular measurements can be found in Ref. [1]. As discussed in the Supporting Information of Ref. [2], in general, the autocorrelation function can be decomposed into two parts:

$$\mathrm{ACF}_{xy}(\tau) = \mathrm{C}_{xy}^2(\tau) + \sigma_{xy}^2 \cdot \delta(\tau)$$

where $\mathrm{C}_{xy}^2(\tau)$ is the autocorrelation of the temporally correlated signal that is likely molecular in origin, $\sigma_{xy}^2$ is the contribution from the temporally uncorrelated noise, and $\delta(t)$ is the Dirac delta function. Thus, when considering time lags $\tau > 0$, all of the temporally uncorrelated noise (*e.g.*, shot noise) will have been removed from an ACF image. As a result, the normalized ACF image for time lags $\tau > 0$ is

$$G_{xy}(\tau > 0) = \frac{C_{xy}^2(\tau)}{C_{xy}^2(0) + \sigma_{xy}^2} = \left( \frac{C_{xy}^2(\tau)}{C_{xy}^2(0)} \right) \cdot \left( \frac{SNR_{xy}^2}{1 + SNR_{xy}^2} \right)$$

where the signal-to-noise ratio, $SNR_{xy} \equiv C_{xy}(0)/\sigma_{xy}$. The first term is the normalized temporal autocorrelation of the only molecular signal (that includes the effects of, *e.g.*, diffusion, blinking, jumps between different emissive states due to conformational dynamics), while the second term depends only on the SNR and controls the contrast of the target molecule in the normalized ACF image. Thus, for nonzero lag times, values of $G_{xy}(\tau)$ approaching 1 indicate the presence of a strong, temporally correlated, molecular signal, whereas values approaching 0 indicate that the pixel intensity is dominated by temporally uncorrelated noise that is unlikely to be a molecule of interest. Target molecules are then located by identifying local maxima pixels in the normalized ACF image, and comparing the $G_{xy}(\tau > 0)$ of each local maximum to an intensity threshold value, $G_0$. A target molecule is determined to be present at a pixel $xy$ when $G_{xy}(\tau > 0) \geq G_0$. We typically use $\tau = 1$ or 2 frames, and $G_0 = 0.15$. For situations where the temporally correlated, molecular signal is extremely strong and long-lived, such that $C_{xy}^2(\tau) \approx C_{xy}^2(0)$, this approach identifies target-molecules with an SNR greater than

$$SNR_{xy}(\tau) \approx \sqrt{\frac{G_0}{1 - G_0}} = \sqrt{\frac{0.15}{0.85}} \approx 0.42,$$

whereas detecting faster and/or weaker molecular signals (*i.e.*, $C_{xy}^2(\tau) < C_{xy}^2(0)$) requires better SNR. Given that most experimental studies would probably exclude single-molecule data with an SNR less than about 2.5, the threshold value of $G_0 = 0.15$ corresponds to detecting target molecules with signals that exhibit a decay of molecular signal correlation less than $C_{xy}^2(\tau)/C_{xy}^2(0) \approx 0.174$. For reference, a stable signal of value $m$ in a time series of length $T$ that photobleaches to a signal value of zero after $fT$ measurements has $C_{xy}^2(\tau) = (fT - \tau)/T \cdot m^2$, and $C_{xy}^2(0) = fT/T \cdot m^2$; so, $C_{xy}^2(\tau)/C_{xy}^2(0) = 1 - \tau/fT$. In a $\tau = 1$ ACF image, any molecule with an SNR of 2.5 and a value of $fT > 1.2$ would pass this threshold; so, target molecules that fluoresce for two or more frames before photobleaching occurs would be detected and localized with this threshold.

*Step 2: Estimating target-associated fluorescence intensity.* After localization, the target-associated fluorescence signal can be estimated by inferring the signal intensity, local background intensity, and point-spread function (PSF) parameters that most likely explain the features of the observed image. This type of inference underlies single-molecule localization microscopy, and the statistical basis of it is well understood [3]. For a two-dimensional, diffraction-limited image with an anisotropic PSF, as many as six parameters may need to be estimated for each target molecule in every frame, creating a substantial computational burden [4]. We reduce this burden by treating the PSF as a known parameter, and using a rapidly converging maximum-likelihood estimation (MLE) algorithm to infer only the target-associated fluorescence intensity and local background intensity for each target molecule in each frame of a TIRF movie [2,5].

For more than ~100 detected photons, which is experimentally reasonable in TIRF-based smF experiments, a Poisson-distributed photon-count distribution can be approximated by a normal distribution. For an isolated target molecule in a local region with a relatively flat amount of background fluorescence, the fluorescence intensity recorded at pixel $xy$ in frame $t$ can be modeled as

$$I_{xy}(t) \sim \mathcal{N}\left( I_{xy} \,\middle|\, \left(B_i(t) + N_i(t) \cdot \Psi_{i,xy}\right), \sigma_i^2(t) \right),$$

where $N_i(t)$ is the target-associated fluorescence intensity for the $i^{th}$ target molecule, $\Psi_{i,xy}$ is the PSF for that molecule integrated over pixel $xy$, $B_i(t)$ is the local background intensity, and $\sigma_i^2(t)$ is the variance of the noise in the local region. Notably, for computational tractability, this assumes that $\sigma_i^2(t)$ is constant and unaffected by, *e.g.*, shot noise differences caused by the varying strength of PSF as it decays away from the central local of the molecule, or pixel-to-pixel differences in amplification, as is the case with scientific complementary metal-oxide semiconductor cameras. In the case of non-isolated molecules, the PSFs of neighboring target molecules can overlap; so, the location parameter of the normal distribution for $I_{xy}(t)$ is actually modeled as

$$B_i(t) + \sum_j N_j(t) \cdot \Psi_{j,xy},$$

where the sum is taken over all localized molecules, including the $i^{th}$ target molecule. Assuming that pixels are independent, the log likelihood probability of the pixels in a region surrounding $i^{th}$ target molecule during frame $t$ is

$$\mathcal{L}_i(t) \approx \sum_{\{xy\}_i} \ln \mathcal{N}\left( I_{xy}(t) \,\middle|\, \left(B_i(t) + \sum_j N_j(t) \cdot \Psi_{j,xy}\right), \sigma_i^2(t) \right),$$

where $\{xy\}_i$ are the pixels in the $\epsilon$-neighborhood analysis window centered on the *i*th target molecule.

The MLE estimate of $N_i(t)$ is obtained by solving $\partial\mathcal{L}_i(t)/\partial N_i(t) = 0$, which yields

$$N_i(t) = \frac{\sum_{\{xy\}_i} \left( \left( I_{xy}(t) - B_i(t) - \sum_{j/i} N_j(t)\Psi_{j,xy} \right) \Psi_{i,xy} \right)}{\sum_{\{xy\}_i} \Psi_{i,xy}^2}.$$

Similarly, the MLE estimate of $B_i(t)$ is obtained by solving $\partial\mathcal{L}_i(t)/\partial B_i(t) = 0$, which yields

$$B_i(t) = \frac{\sum_{\{xy\}_i} \left( I_{xy}(t) - \sum_j N_j(t)\Psi_{j,xy} \right)}{\sum_{\{xy\}_i} 1}.$$

The MLE estimates of $N_i(t)$ and $B_i(t)$ are coupled, but optimal values can be found by iterating successive updates of these two parameters using the formulas given above. Reasonable initializations are

$$B_i(t) \approx \frac{\sum_{\{xy\}_i} I_{xy}(t)}{\sum_{\{xy\}_i} 1},$$

and

$$N_i(t) \approx I_{xy=i}(t) - B_i(t),$$

where $xy = i$ denotes the pixel containing the center of the *i*th target molecule. Because the local background is estimated separately for each frame, this approach is particularly useful for analyte-delivery experiments in which the background intensity varies over time.

For computational efficiency, we perform these updates using a square $\epsilon$-neighborhood of the *i*th target molecule with $\epsilon = 11$ (*i.e.*, using the 121 pixels in an 11 × 11-pixel square analysis window centered on each target molecule). This window is sufficient to encompass the PSFs produced by most single-molecule fluorescence microscopes. The functional form of $\Psi_{xy}$ is typically taken as an isotropic two-dimensional Gaussian integrated over the area of each pixel [4] with either a theoretically determined width or one fitted from experimental data. In the latter case, it is worth noting that a two-point ACF image is proportional to the square of the PSF; so, the width of a Gaussian $\Psi_{xy}$ in an ACF image is $1/\sqrt{2}$ times that of the diffraction-limited width.

**4. Detailed Slide Surface Functionalization Protocol**

The protocol described below yields exceptionally high-quality, functionalized quartz flow cells. Many steps, such as slide-cleaning and flow cell assembly steps, were adapted from conventional surface-functionalization protocols [6,7]. Certain steps in this protocol involve potential hazards, including drilling through glass, handling large volumes of caustic and flammable liquids, and working with open flames. Researchers must wear appropriate personal protective equipment (PPE) at all times, including safety glasses, flame-resistant clothing, and chemical-resistant gloves. Prior to beginning, familiarize yourself with the location of all relevant fire-safety equipment, including safety showers, eyewash stations, and fire blankets. In our experience, quartz flow cells properly functionalized using this protocol deteriorate in quality over time but remain suitable for micromolar concentration smF imaging experiments for at least two to three days when stored in a covered slide dish at room temperature between preparation and use. Certain experimental systems may see degraded quality after four or more days. We prioritize performing experiments on the first two days after functionalization.

Three to five natural quartz microscope slides (1” × 3” × 1 mm; G. Finkenbeiner) are prepared in a batch. Using a diamond burr drill bit with a 0.03-inch head diameter (McMaster-Carr; catalog no. 4490A75), drill five pairs of inlet/outlet holes along the long edges of each quartz microscope slide. All holes should be confined to the central 1” region of the 3”-long side of the slide. For each pair of holes, the holes are positioned approximately 3 mm from the nearest long edge, and spaced approximately 5 mm apart across the length of the slide. Each pair serves as the inlet and outlet ports for one of five flow cells that span the 1” side of the slide. After drilling, thoroughly rinse each slide with ultrapure water, then keep it submerged in ultrapure water in a glass, Coplin, slide-staining jar (BRAND GMBH; catalog no. 472800). After all holes have been drilled, sonicate the slides in an ultrasonic water bath at room temperature for approximately 15 s to dislodge residual glass debris. Decant the water, then rinse the slide-staining rack with ultrapure water several times to remove any glass debris.

Fill the slide-staining rack with enough 1% *w/v* Alconox solution to completely immerse the drilled slides. Incubate the slides in the Alconox solution for at least 2 hours at room temperature; incubations of up to 24 hours did not impair subsequent surface functionalization. One by one, remove each slide and, while its surfaces remain coated with detergent solution, gently scrub both faces with a freshly gloved fingertip for ~30 s using nitrile gloves. Rinse the slide with ultrapure water flowing from a dispenser spigot until no visible detergent remains, and transfer it to a clean, glass slide-staining rack filled with fresh ultrapure water. After all slides have been scrubbed and rinsed, replace the water in the slide-staining rack with enough freshly prepared 1 M potassium hydroxide (KOH) to immerse the slides completely (*Caution: 1 M KOH is a caustic solution capable of producing chemical burns; wear appropriate PPE; avoid contact with skin and eyes*). Incubate for 2-3 hours at room temperature. Longer KOH incubation times have not been systematically evaluated.

After incubation in KOH is complete, transfer the slides from the glass slide-staining rack to a plastic slide-staining tray (BRAND GMBH; catalog no. 474305). Place the same number of No. 1.5 borosilicate coverslips (24 mm x 30 mm; VWR) in an alumina coverslip staining rack (*n.b.*, to our knowledge, such products have been discontinued; a plastic coverslip-staining rack, such as the Diversified Biotech WSDR-1000 is a suitable replacement). Add two extra coverslips in case some are dropped or cracked during subsequent steps.

Place the slide-staining tray and the coverslip-staining rack inside polypropylene copolymer container of 500 mL and 250 mL, respectively (Nalgene; catalog no. 2118-0016). Add sufficient absolute ethanol to each container to completely immerse the slides and coverslips. Use an ultrasonic water bath to sonicate the containers at room temperature for 20 minutes. Remove the ethanol, add sufficient 1 M KOH to completely immerse the slides or coverslips, and sonicate at room temperature for an additional 30 min. Rinse each slide and coverslip thoroughly with ultrapure water flowing from a dispenser spigot to remove residual KOH, then dry with nitrogen or argon passed through a 0.22-µm filter. Allow the dried slides and coverslips to sit for 15 minutes in a clean, dust-protected environment (*e.g.*, a drawer) to permit residual moisture to evaporate before flame cleaning.

Flame-clean the slides and coverslips to remove residual organic and fluorescent contaminants (*Caution: This procedure uses an open flame. Wear flame-resistant PPE, properly store away all flammable solvents, and tie back long hair before operating the torch*). Perform this step only after confirming that the slide and coverslip surfaces are completely dry. Hold each slide with forceps approximately 1 cm from the nozzle of a propane torch and pass each face slowly through the outer region of the flame four times, ensuring that the flame traverses the drilled port and intended flow-channel regions. Each pass should require approximately 10 s. Transfer the slide to a clean, glass slide-staining jar and allow it to cool to room temperature before returning it to the plastic slide-staining tray. Before flame-cleaning the coverslips, inspect them carefully and confirm that no water droplets remain, as residual water can cause uneven heating that will deform the glass. Hold each coverslip with wafer tweezers (Excelta; catalog no. 35A-SA), pass each face rapidly through the outer region of the flame three times, and transfer it to a clean, alumina coverslip-staining rack (or another non-plastic container to avoid melting the plastic) to cool until it reaches room temperature.

Methoxy-terminated PEG-silane (mPEG-silane; 5 kDa; Laysan Bio; catalog no. MPEG-SIL-5000-5GR) and biotin-terminated PEG-silane (biotin-PEG-silane; 5 kDa; Laysan Bio; catalog no. BIOTIN-PEG-SIL-5000-1GR) were divided on receipt into 50-mg and 20-mg aliquots, respectively, in separate microcentrifuge tubes. Aliquots were stored at –20 °C in a nitrogen-flushed desiccator until use. Once the flame-cleaned slides and coverslips have cooled, the following steps are performed as quickly as possible.

Dissolve 20 mg biotin-PEG-silane in 1 mL absolute ethanol to prepare a 20 mg $mL^{-1}$ stock, and vortex thoroughly to facilitate dissolution. Separately, dissolve 50 mg mPEG-silane in 450 $\mu$L of 2% *v/v* aqueous acetic acid, prepared by combining 1 mL glacial acetic acid with 49 mL ultrapure water. Bubbles may appear in this solution. Briefly centrifuge the tube to collect the liquid and foam from the walls, and confirm visually that no undissolved material remains.

Dilute 5 $\mu$L of the 20 mg $mL^{-1}$ biotin-PEG-silane stock solution to a final volume of 1 mL with 2% *v/v* aqueous acetic acid. Quickly, add 50 $\mu$L of this diluted solution of biotin-PEG-silane, containing 5 $\mu$g, to the 50-mg mPEG-silane solution. This produces 500 $\mu$L of PEG-silane solution with a 10,000:1 mPEG-silane:biotin-PEG-silane mass ratio. Quickly, add 500 $\mu$L of 1.5 M aqueous magnesium sulfate ($MgSO_4$), prepared fresh from a 2.5 M aqueous stock. The resulting 1-mL mixture contains approximately 5% *w/v* total PEG-silane, 750 mM $MgSO_4$, and 1% *v/v* acetic acid. Briefly, vortex the mixture to ensure mixing. The solution should become cloudy as phase separation begins. Centrifuge at >20,000 × g for 2.5 minutes at room temperature. An aqueous biphasic system should form; do not continue if two distinct layers are not seen, as phase separation did not occur. In such a case, a higher concentration of $MgSO_4$ is required to reach the cloud point. Stop the current preparation and empirically determine the $MgSO_4$ concentration required to reach the cloud point (*e.g.*, using 50 mM increments of $MgSO_4$). Once determined, discard all PEG-silane solutions, and prepare a fresh solution at the newly determined $MgSO_4$ concentration for the subsequent steps.

Place each microscope slide on a clean, level surface. Ensure the side through which holes were drilled faces downward and the blown-out side faces upward. Apply 70 $\mu$L of the lower, PEG-depleted phase to the center of the upward-facing surface between the drilled holes. Carefully lower a cleaned coverslip onto the solution to form a slide-solution-coverslip sandwich, ensuring that the solution spreads across all five intended flow-cell regions without visible air bubbles. If air bubbles are present, gently tap the edges of the coverslip with a clean, plastic pipette tip to release them. Keep the coverslip aligned with the slide throughout functionalization; significant displacement can leave portions of the intended flow-cell regions unfunctionalized, which reduces the number of usable flow cells.

Place the assembled slide-solution-coverslip sandwiches on the rack from a 10-$\mu$L pipette-tip box positioned inside a closed 1000-$\mu$L pipette-tip box containing an approximately 1-cm-deep layer of water. Ensure that the assemblies remain above the water and do not contact it. This arrangement serves as a humidity chamber. Place the container in a closed drawer where the assemblies can incubate undisturbed for 20-22 hours at room temperature.

After functionalization, use a carbide-tipped pencil to mark an edge of each slide and coverslip so that the functionalized face can be identified without scratching the intended imaging regions. Carefully separate each slide-coverslip pair, rinse both components thoroughly with fresh ultrapure water flowing from a dispenser spigot and dry them using 0.22-$\mu$m-filtered nitrogen or argon. Place narrow strips of double-sided tape (3M; Scotch 665) on the functionalized side of each microscope slide to separate each inlet-outlet hole pair and define five parallel flow channels. Ensure a strip is also placed above the first, and below the last inlet-outlet hole pair (six strips total). Position the corresponding coverslip on the tape with its functionalized face oriented toward the functionalized slide surface. Use a plastic pipette tip to gently apply force to the coverslips in order to ensure adherence (*n.b.*, only press above the tape strips or the coverslip may crack). Seal the perimeter and ends of each flow cell with epoxy (5 Minute Epoxy; Devcon), leaving the drilled inlet and outlet ports unobstructed. Allow the epoxy to cure for at least one hour. Cut 200-$\mu$L pipette tips at the first notch, approximately 0.8 cm from the top, and affix the cut end to each inlet port with epoxy to create a solution reservoir. Do not block the inlet/outlet holes. During solution delivery, connect the corresponding outlet hole to a syringe pump to draw solutions loaded into the inlet reservoir through the flow cell.
